# Small allelic variants are a source of ancestral bias in structural variant breakpoint placement

**DOI:** 10.1101/2023.06.25.546295

**Authors:** Peter A. Audano, Christine R. Beck

**Affiliations:** The Jackson Laboratory for Genomic Medicine, Farmington, CT, USA; Department of Genetics and Genome Sciences, Institute for Systems Genomics, University of Connecticut Health Center, Farmington, CT, USA

## Abstract

High-quality genome assemblies and sophisticated algorithms have increased sensitivity for a wide range of variant types, and breakpoint accuracy for structural variants (SVs, ≥ 50 bp) has improved to near basepair precision. Despite these advances, many SVs in unique regions of the genome are subject to systematic bias that affects breakpoint location. This ambiguity leads to less accurate variant comparisons across samples, and it obscures true breakpoint features needed for mechanistic inferences. To understand why SVs are not consistently placed, we re-analyzed 64 phased haplotypes constructed from long-read assemblies released by the Human Genome Structural Variation Consortium (HGSVC). We identified variable breakpoints for 882 SV insertions and 180 SV deletions not anchored in tandem repeats (TRs) or segmental duplications (SDs). While this is unexpectedly high for genome assemblies in unique loci, we find read-based callsets from the same sequencing data yielded 1,566 insertions and 986 deletions with inconsistent breakpoints also not anchored in TRs or SDs. When we investigated causes for breakpoint inaccuracy, we found sequence and assembly errors had minimal impact, but we observed a strong effect of ancestry. We confirmed that polymorphic mismatches and small indels are enriched at shifted breakpoints and that these polymorphisms are generally lost when breakpoints shift. Long tracts of homology, such as SVs mediated by transposable elements, increase the likelihood of imprecise SV calls and the distance they are shifted. Tandem Duplication (TD) breakpoints are the most heavily affected SV class with 14% of TDs placed at different locations across haplotypes. While graph genome methods normalize SV calls across many samples, the resulting breakpoints are sometimes incorrect, highlighting a need to tune graph methods for breakpoint accuracy. The breakpoint inconsistencies we characterize collectively affect ∼5% of the SVs called in a human genome and underscore a need for algorithm development to improve SV databases, mitigate the impact of ancestry on breakpoint placement, and increase the value of callsets for investigating mutational processes.

## Introduction

The human reference genome (International Human Genome Sequencing Consortium, 2001; Schneider et al., 2017) hosts annotations including genes (Frankish et al., 2021; O’Leary et al., 2016), regulatory regions (Encode Project Consortium, 2012; Encode Project Consortium et al., 2020), and repeats (Bailey et al., 2002; Benson, 1999; Smit, 2013-2015), and it has become a universal coordinate system for describing genetic alterations across populations (Abel et al., 2020; Audano et al., 2019; Beyter et al., 2021; Collins et al., 2020; Ebert et al., 2021; International HapMap et al., 2007; Karczewski et al., 2020; Sudmant et al., 2015; The 1000 Genomes Project Consortium, 2015) and diseases (ICGC TCGA Pan-Cancer Analysis of Whole Genomes Consortium, 2020; Taliun et al., 2021; Turner et al., 2017). New high-quality references are emerging for humans (Nurk et al., 2022) and a growing number of other species (Alonge et al., 2020; Ferraj et al., 2023; Jebb et al., 2020; Li et al., 2023; Mao et al., 2021; Mouse Genome Sequencing Consortium et al., 2002), which play fundamental roles in modern genomics.

Variant discovery is largely based on aligning reads or assemblies to a reference genome. This is used to identify single nucleotide variants (SNVs), small insertions and deletions (indels), and structural variants (SVs) including insertions and deletions ≥ 50 bp, inversions, complex rearrangements, and chromosomal translocations. Imprecise SV breakpoints affect comparisons across samples, and while new methods are improving comparisons (Ebert *et al*., 2021; English et al., 2022; Kirsche et al., 2021), error-free merging across many haplotypes has not yet been attained. Additionally, breakpoint features such as microhomology and nearby variants in-*cis* are important signatures for predicting mechanisms of formation (Beck et al., 2015; Carvalho and Lupski, 2016; Carvalho et al., 2011; Vogt et al., 2014). Repetitive sequences often mediate SVs and can make the determination of precise breakpoints challenging.

Recent advances in sequencing technology are now generating longer and more accurate reads capable of reaching into repetitive structures and spanning larger SVs. As a result, many new SV loci have been discovered, and SV yield per sample has increased from less than 10,000 SVs per genome to more than 25,000 (Audano *et al*., 2019; Chaisson et al., 2015; Ebert *et al*., 2021). Moreover, long-reads routinely reveal the full sequence of SVs, which was not previously attainable. In more recent years, long-read phased assemblies have become a critical component for producing complete and accurate variant callsets (Chaisson et al., 2019; Ebert *et al*., 2021; Garg et al., 2021; Liao et al., 2023). These advances enable more complete transposable element (TE) analysis, improve genotyping in short-read samples, and support new biological insights (Ebert *et al*., 2021; Ebler et al., 2022; Rozowsky et al., 2023).

Modern references are a single theoretical human haplotype creating alignment biases when reads are mapped to non-reference alleles, which can be difficult to mitigate (Brandt et al., 2015; Degner et al., 2009; Eizenga et al., 2020). To support mapping and variant calling across diverse genomes, the Human Pangenome Reference Consortium (HPRC) is developing graph-based references encompassing many haplotypes simultaneously (Liao *et al*., 2023). While in-graph haplotypes can be directly detected, variants absent from the graph reference still rely on calling differences between the sample and a graph path. Therefore, challenges with linear reference analyses will ultimately translate to graphs, especially for rare and somatic events often associated with disease (Nattestad et al., 2018; Rausch et al., 2023; Sakamoto et al., 2020; Vogt *et al*., 2014; Wahlster et al., 2021). However, in-graph SVs are represented as a unique "bubble" in graph space with a common breakpoint across samples, and so ambiguity with merging across independent haplotypes may be eliminated, although breakpoint accuracy has not been assessed.

While contiguous high-accuracy assemblies are becoming routine, we find that SV breakpoints are still inconsistently placed across phased haplotypes, and many breakpoints do not represent the true site of rearrangements, potentially impeding downstream analyses. To quantify the effect on modern long-read variant discovery approaches, we re-analyze a recent callset from 64 phased haplotypes recently released by the Human Genome Reference Consortium (HGSVC) (Ebert *et al*., 2021). With pangenomes recently released by the Human Pangenome Reference Consortium (HPRC) (Liao *et al*., 2023), we identify discordance between linear- and graph–based-reference approaches. We determine reasons why breakpoints can differ between assemblies and suggest approaches for improving both mechanistic inference and variant comparisons across samples.

## Results

### Breakpoint offsets are prevalent in long-read SV callsets

We examined breakpoint placement for SVs across 64 phased haplotypes derived from 32 diverse samples released by the HGSVC (Ebert *et al*., 2021). In that study, variants were called independently on each assembled haplotype against the GRCh38 reference using minimap2 (Li, 2018) and merged to a multi-haplotype, nonredundant callset. For each pair of haplotypes (2,016 combinations of 64 haplotypes), we find an average of 20% of insertions and 15% of deletions have different breakpoints between the pair. When SVs anchored in tandem repeats (TRs) and segmental duplications (SDs) are excluded (Methods), we find 4.4% of insertions and 1.7% of deletions on average have different breakpoints (**Table S1**). In this paper, we refer to "unique loci" as regions outside TRs and SDs where large and highly repetitive structures often produce ambiguous alignments. We include transposons in our unique loci because the 64 assemblies we are investigating (average n50 > 19.5 Mbp) are capable of spanning full-length human TEs, which cluster around 300 bp and 6 kbp.

Inconsistent breakpoints in unique regions affect a small number of variants per haplotype pair, but the effect across multiple haplotypes and samples is greater. In the merged callset across all 64 haplotypes, we find 5.9% insertions and 3.1% deletions in unique loci disagree on breakpoint location (**Table 1**). While many of these differences are small, insertions vary by a median of 2.2 bp and deletions by 4.9 bp resulting in a non-trivial effect on SV representation (**Fig 1A**, **Table 1**).

**Figure 1:**
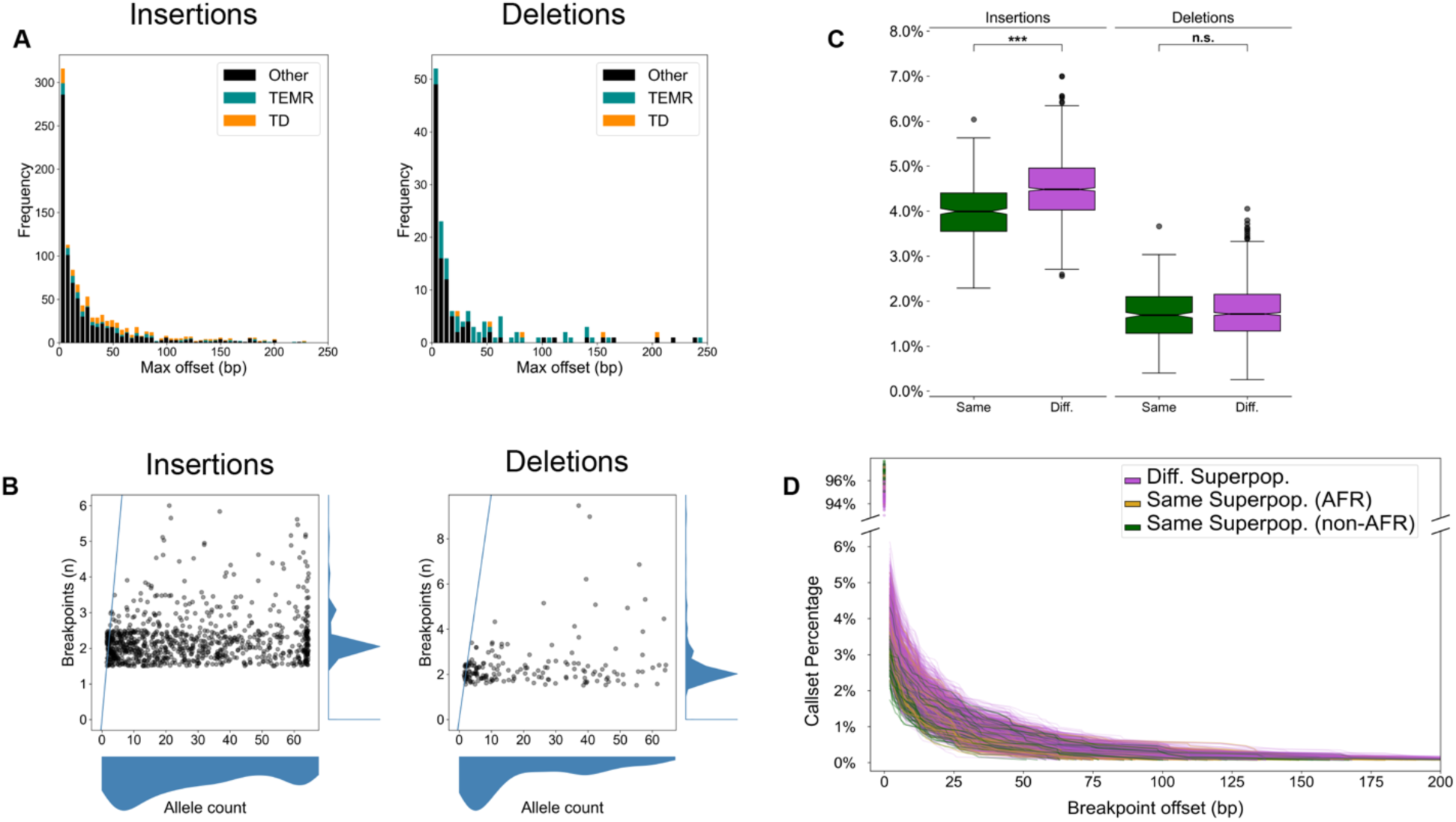
Breakpoint offsets reflect population structure. (A) Maximum offset distances for variants in the merged callset. For SVs identified in more than one haplotype, the maximum offset is the difference betwee and right-most breakpoints. TEMR: Transposable element mediated rearrangements. TD: Tandem duplications. **(B)** The number of unique breakpoints for each variant (vertical axis) does not scale with the number of haplotypes (horizontal axis). A blue line represents the x = y diagonal. Scatterplot points were jittered in each axis uniformly from -0.5 to 0.5 to show density. **(C)** For any pair of haplotypes, the proportion of offset SVs is stratified by same superpopulation (green) or different superpopulation (violet). The difference in means is significant for both insertions and deletions (Student’s T-test of means), but a greater effect is seen for insertions. Notches indicate a 95% confidence interval around the median. n.s: Not significant, *: 1e-3 < p ≤ 1e-2, **: 1e-4 < p ≤ 1e-3, ***: p ≤ 1e-4. **(D)** A cumulative distribution of breakpoint offset for all haplotype pairs. Most variants in both haplotypes share the same breakpoint (upper y-axis). For variants with at least 1 bp offset (lower y-axis), the cumulative proportion of matched calls decreases with increasing breakpoint distance. When both samples come from a different superpopulation (violet), larger differences between breakpoints are observed than when haplotypes come from the same superpopulation (green). When both haplotypes come from African samples (gold), breakpoint distances are elevated, but to a lesser extent than different ancestral backgrounds.

**Table 1:**
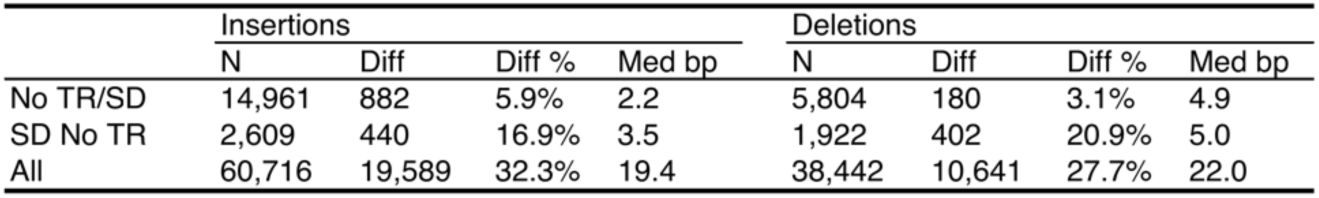
Summary of differential breakpoints in the merged callset. Variants in unique regions (No TR/SD) have the fewest breakpoint offsets, which grows as SVs in more complex loci are included. TR: Tandem Repeat, SD: Segmental Duplication, N: Number of variants, Diff: Variants with different offsets in at least one haplotype, Med: Median breakpoint offset.

|  | Insertions |  |  |  | Deletions |  |  |  |
| --- | --- | --- | --- | --- | --- | --- | --- | --- |
|  | N | Diff | Diff % | Med bp | N | Diff | Diff % | Med bp |
| No TR/SD | 14,961 | 882 | 5.9% | 2.2 | 5,804 | 180 | 3.1% | 4.9 |
| SD No TR | 2,609 | 440 | 16.9% | 3.5 | 1,922 | 402 | 20.9% | 5.0 |
| All | 60,716 | 19,589 | 32.3% | 19.4 | 38,442 | 10,641 | 27.7% | 22.0 |

Finally, the number of distinct breakpoints for each variant does not scale linearly with the number of haplotypes harboring the SV (AC: allele count) (**Fig 1B**). This suggests that variant breakpoints are placed consistently across many haplotypes, but are affected for a subset of haplotypes.

### Diversity is the main driver of differential breakpoint placement

To examine whether random sequence errors may affect SV quality, we compared CLR (21 genomes) with HiFi (11 genomes) in the HGSVC callset. We find a marginally significant enrichment for differential breakpoints in CLR vs HiFi for insertions (4.40% vs 4.29%, *p* = 0.025, Student’s t-test) and no enrichment for deletions (1.75% vs 1.77%, *p* = 0.52, Student’s t-test), which we confirmed with permutation tests (*p* = 0.012 insertions, *p* = 0.74 deletions, 100,000 permutations).

We next asked whether sequence variation affected placement. To investigate this, we stratified the callset by ancestry across 64 HGSVC haplotypes derived from all five 1000 Genomes ancestral superpopulations composed of African, admixed American, East Asian, European, and South Asian ancestry. We observed that variant breakpoints differed more often when a pair of samples were derived from different superpopulations for insertions (4.49% vs 3.99%, *p* = 2.44×10^-40^, Welch’s t-test, Cohen’s D = 0.73) (**Fig 1C**). Deletions were also increased, but the effect did not reach significance (1.76% vs 1.71%, *p* = 0.069, Welch’s t-test) (**Fig 1C**).

Furthermore, there is a noticeable increase in offset distance when haplotypes are derived from different ancestral backgrounds (**Fig 1D**); we confirmed these results with permutation tests (*p* < 1×10^-5^ insertions, *p* = 0.041 deletions, 10,000 permutations). Our results suggest that allelic polymorphisms in or near SVs are a driver of breakpoint differences, which reveals a source of ancestral bias in modern SV callsets.

### Breakpoint offsets are more prevalent with TE-mediated SVs

Transposable elements (TEs) create tracts of homology throughout the genome resulting in TE-mediated rearrangements (TEMRs) (Balachandran et al., 2022; Han et al., 2008; Sen et al., 2006). TEs from the same family have highly similar sequences, and so there are many choices for breakpoint placement along TE copies (**Fig. 2A**). While TEs may provide the homology necessary for duplications and deletions by non-allelic homologous recombination (NAHR), most exhibit only short tracts of breakpoint homology and appear to be mediated by other repair processes (Balachandran *et al*., 2022). Therefore, accurately placing SV breakpoints within TEMRs is essential for understanding the mutational mechanisms underlying their formation (Morales et al., 2015).

**Figure 2:**
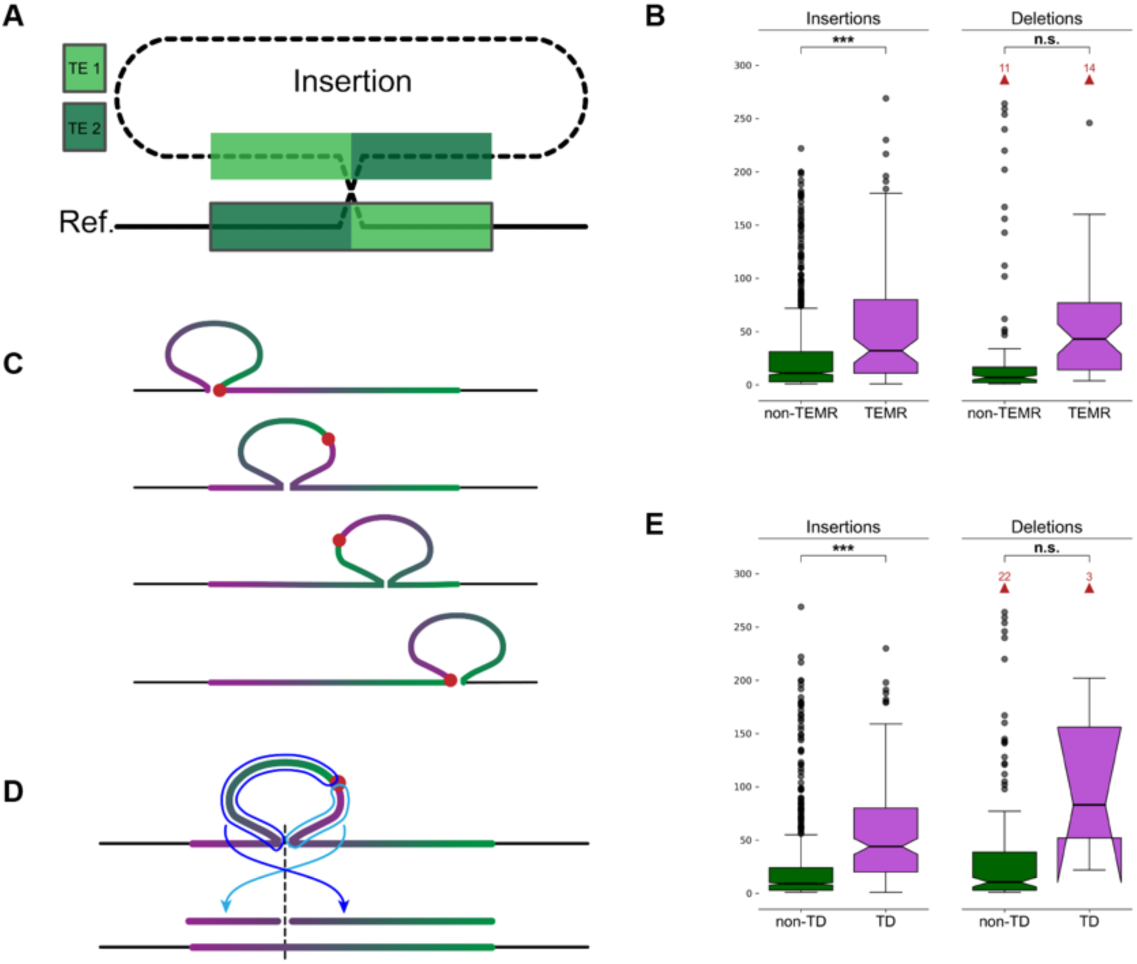
Breakpoints shift through homology and alter SV representation. (A) A non-reference sequence with chimeric TE copies (light green and dark green) at the breakpoints. **(B)** The maximum distance between differential breakpoints across haplotypes is greater in TEMR variants with a significant difference seen for insertions. **(C)** The sequence of a tandem duplication is rotated as the alignment shifts along the reference copy. The true breakpoint (red dot) is located at the insertion breakpoint if the duplicate copy is aligned to the left or right end of the reference copy (top and bottom examples), and it is embedded within the insertion otherwise creating a chimera of duplication copies. **(D)** Mapping a rotated TD to the reference occurs in two pieces separated at the TD breakpoint (light and dark blue arrows) where each piece maps to one side of the reference insertion site (dashed line). Alignment programs often miss one or both fragments. **(E)** Maximum breakpoint offsets in TDs and non-TD SVs. Horns extending downward on the TD deletions indicates that the 95% confidence interval for the median extends below the bottom quartile. (**B**, **E**) *p*-values are generated from T-tests of the mean. Notches indicate a 95% confidence interval around the median. Red arrows and numbers indicate the number of outlier points above the horizontal axis maximum. n.s.: Not significant, *: 1e-3 < *p* ≤ 1e-2, **: 1e-4 < *p* ≤ 1e-3, ***: *p* ≤ 1e-4.

In the merged HGSVC SV callset, we find 112 SV insertions and 119 SV deletions with differential breakpoints in unique loci are likely TEMRs (8.5% and 20.4% of differential variants, respectively) (Methods). We find TEMR insertions were significantly enriched for offset breakpoints (odds ratio (OR) = 4.18, *p* = 3.17×10^-25^, Fisher’s exact test (FET)), as were TEMR deletions (OR = 3.20, *p* = 1.55×10^-11^, FET). TE homology is also responsible for larger distances between breakpoints across haplotypes for insertions (15.17 vs 2.50 bp, *p* = 1.45×10^-8^, Welch’s T-test), but the breakpoint difference was did not reach significance for deletions (46.71 vs 10.93 bp, *p* = 0.065, Welch’s T-test) (**Fig 2B**).

### Tandem duplications are heavily affected by differential breakpoints

Tandem duplications (TDs) are a common SV type where a duplicate copy is inserted adjacent to its template. TDs may be driven by existing homology, such as NAHR, or occur in regions with little to no homology (Arlt et al., 2009; Lee et al., 2007; Li et al., 2020; Menghi et al., 2016; Willis et al., 2017). With short reads, TDs are detected by elevated copy number of the duplicated sequence combined with paired-end evidence at the duplication breakpoint, revealing the duplicated reference region (Alkan et al., 2011). However, long-read callers often call TDs as insertions, especially when assemblies are used. A TD is highly homologous with itself posing a significant problem for alignment algorithms because there are many valid choices for the breakpoint placement. If the breakpoint is not placed on one end of a reference copy, the insertion sequence contains a chimera of both duplication copies and the true breakpoint is embedded somewhere inside the insertion sequence (**Fig 2C**). Annotating a TD should be as easy as re-aligning the insertion sequence to the reference and determining if it maps adjacent to the insertion breakpoint, however, rotated TDs align in two separate fragments (**Fig. 2D**), and current alignment programs often miss one or both fragments. To better annotate SVs as TDs, we re-aligned SV sequences with BLAST (Altschul et al., 1990) (Methods). We found 1,843 SV insertions were TDs, of which 261 (14.2%) were shifted and rotated leading to the duplication mapping to two separate BLAST records on each side of the insertion site. We found 17 reference TDs with a deleted copy, of which 8 (47.1%) were shifted and rotated mapping to both sides of the deletion SV.

We find that TD insertions are more likely to have differential breakpoints (OR = 0.55, *p* = 1.45×10^-9^, FET), but the effect on TD deletions is small (OR = 0.05, *p* = 3.14×10^-7^, FET). Because homology runs across the full length of a TD, we observe greater average offset distances for insertions (9.37 bp TD vs 2.20 bp non-TD, *p* = 1.07×10^-13^, Welch’s t-test, Cohen’s D = 0.45). A large increase in distance for deletions failed to reach significance (741.9 bp TD vs 12.8 bp non-TD, *p* = 0.19, Welch’s t-test, Cohen’s D = 2.23) (**Fig. 2E**). These results appear to suggest that TD deletions are highly susceptible to breakpoint shifts, however, the low number of these events and the large range of offsets across all deletions make these observations difficult to validate.

### Small polymorphisms surround offset SV breakpoints

Differential breakpoints occur in regions with tracts of homology, such as TEMRs and TDs. In cases of perfect homology, the actual breakpoint could have occurred anywhere in the homologous region. By convention, many aligners, such as minimap2 (Li, 2018), push breakpoints to the left yielding more consistent variant calls across haplotypes. As we have observed, variation in breakpoint placement increases when haplotypes are derived from different ancestral superpopulations, therefore, we reasoned that small allelic polymorphisms around SV breakpoints might influence alignments.

To identify small polymorphisms at SV breakpoints that might influence alignments, we extracted the offset region around breakpoints from each haplotype assembly and compared them (**Fig 3A**) (Methods). For SV insertions in unique regions with shifted breakpoints, we find on average 5.0 small variants on the left breakpoint vs 5.2 on the right breakpoint (*p* = 1.62×10^-10^, Welch’s t-test) (**Fig 3B**).

**Figure 3:**
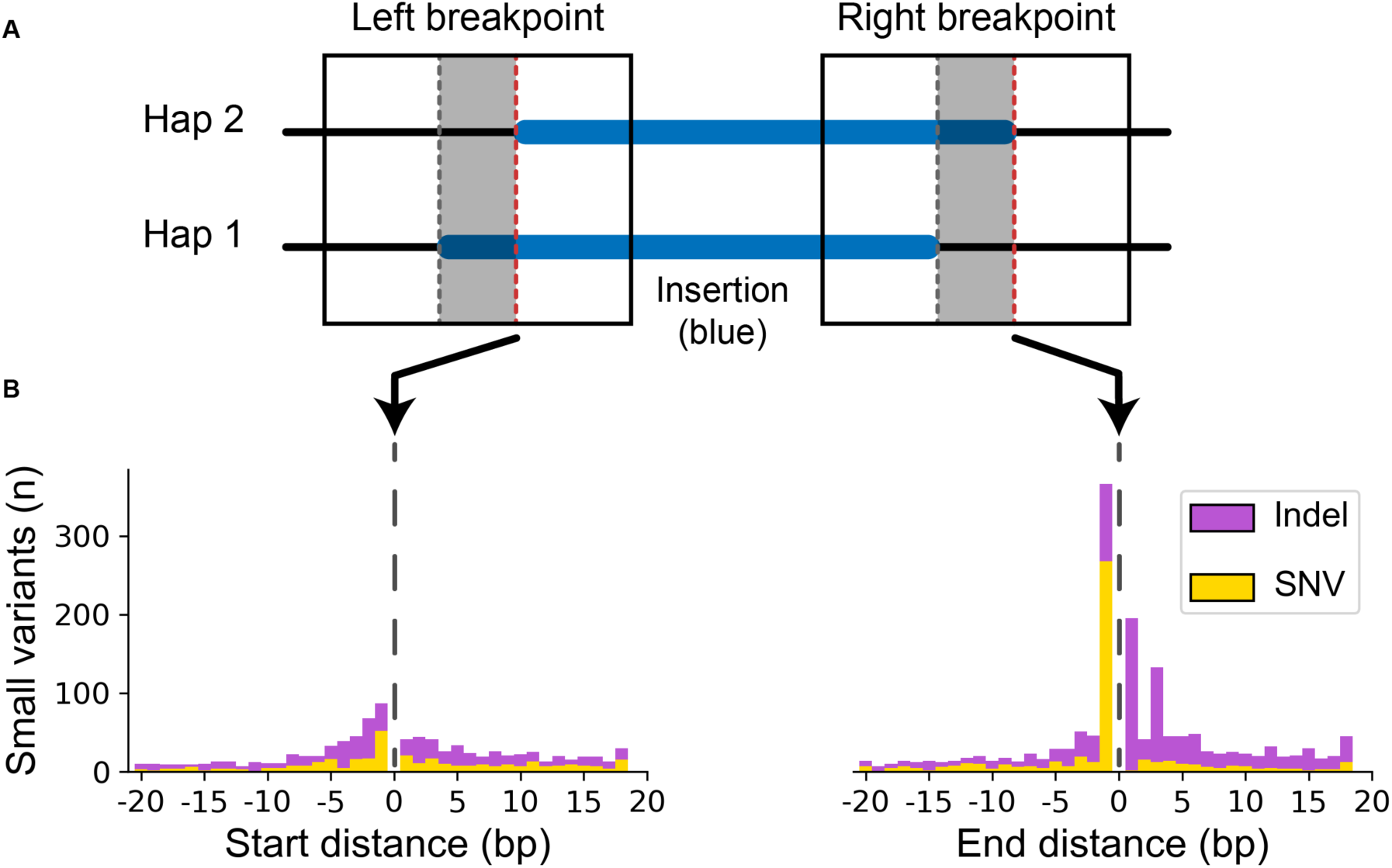
Small variants accumulate at differential SV breakpoints. (A) SV insertions with different breakpoints (blue) in each haplotype pair were retrieved. Sequence around the left and right breakpoints was extracted (solid box) for both haplotypes including the differential locus (gray area between dashed lines) and 50 bp upstream and downstream. The red dashed line marks the start and end position of the right-shifted variant. **(B)** Small variants accumulate at the upstream and downstream edges of the right-shifted variants where 0 is the red line in part (A).

When polymorphisms occur in homologous regions, it creates differences between the haplotype and the reference sequence, which penalizes the alignment score for the haplotype more diverged from the reference. If the variant can be shifted across a homologous region, a better alignment score may be achieved by moving the breakpoint such that small polymorphisms are pushed into the unaligned insertion sequence. As a consequence, we observe a large peak of small polymorphisms on the upstream breakpoint shifted 1 bp inside the insertion (**Fig 3B**). Not only does this drive breakpoint disagreements, but these small polymorphisms near SVs are systematically removed from variant callsets.

### Breakpoint homology annotations change with breakpoint placement

SVs are often mediated by large tracts of homology during their formation through NAHR requiring more than 100 bp of perfect homology, double-strand break repair pathways mediated by tracts of microhomology from 1 to 10 bp, non-homologous end joining (NHEJ) requiring no breakpoint homology, and alternative end-joining (alt-EJ) requiring little or no microhomology (reviewed in Carvalho and Lupski (2016) and Iliakis et al. (2015)). Mobile element insertions (MEIs) create homology in the form of target-site duplications (TSDs), which is an important annotation for distinguishing true MEI polymorphisms from other SVs containing MEI sequence (Ebert *et al*., 2021; Zhou et al., 2020). Accurately detecting breakpoint homology is an important predictor of SV mechanism and a useful quality metric for SV callsets, however, the effect of differential breakpoints on microhomology has not been investigated.

We used a recent PAV addition to estimate microhomology for all SV breakpoints (Balachandran *et al*., 2022) (Methods). For each SV, we find that the number of breakpoints increases the number of different microhomology calls for insertions (ρ = 0.72, *p* < 1×10^-100^, Spearman rank-order correlation (Spearman)) and deletions (ρ = 0.87, *p* < 1×10^-100^, Spearman) indicating that breakpoint changes affect homology annotations in almost all cases (**Fig. S1**). For insertions with consistent breakpoints (n = 6,855), microhomology annotations varied by 2.16 bp on average, which rises to 21.91 bp on average with inconsistent breakpoints (n = 725) (*p* = 9.46×10^-15^, Welch’s t-test, Cohen’s D = 0.43). We see a similar effect on deletion microhomology, which varies by 0.01 bp across haplotypes with consistent breakpoints (n = 3,399) and rises to 19.27 bp across haplotypes with inconsistent breakpoints (n = 172) (*p* = 1.01×10^-16^, Welch’s t-test, Cohen’s D = 1.77) (**Fig. S2**).

As a result of imprecise breakpoints, actual breakpoint homologies necessitate manual reconstruction, which is tedious task and cannot easily scale with modern whole-genome analyses. Therefore, precise mechanisms are difficult to annotate at scale. For example, while SVs mediated by mobile elements with at least 85% identity are generally thought to be mediated by non-allelic homologous recombination (NAHR) (Lam et al., 2010), a closer examination of breakpoints using modern long-read data shows that at least 20% have breakpoint features inconsistent with NAHR (Balachandran *et al*., 2022).

### Read-based approaches have less consistent breakpoints

We examined the effects of offset breakpoints from aligned reads using PBSV. In our callset, 11,906 SV insertions and 5,501 SV deletions in unique loci were callable across the HGSVC samples. We find that 13% of insertions and 18% of deletions are offset across samples when called from read alignments, which is higher than 6% insertions and 3% deletions we observe from assembly-based callsets for the same SVs. We hypothesize that assemblies are more consistent because a single polished representation of the region is aligned where individual reads may be subject to more systematic bias, for example read errors and SVs on the edges of individual reads.

While short-read callers typically rely on read alignments, some produce breakpoint assemblies and may improve breakpoint accuracy. A recent study of TEMRs (Balachandran *et al*., 2022) finds that MANTA (Chen et al., 2016) places SVs more accurately than other short-read callers, which may be a result of breakpoint assemblies MANTA performs.

### Pangenomes

Pangenome graphs are constructed from multiple haplotypes and can be used to negate differences in alignments. The Pangenome Graph Builder (PGGB) (Garrison et al., 2023) constructs graphs from multiple haplotypes simultaneously, and the Minigraph-Cactus (MC) approach iteratively adds haplotypes to a graph (Hickey et al., 2022). Both were featured in the recent pangenome drafts constructed from 94 phased assemblies derived from 47 diverse samples recently released by the Human Pangenome Reference Consortium (HPRC) (Liao *et al*., 2023).

Across unique loci, we identified all SVs that were present in more than one haplotype and matched an SV identified by MC (4,851 insertions, 3,240 deletions). We find that the MC breakpoint offset is greater than all the HGSVC offsets for 69% of SV insertions and 41% of SV deletions. For PGGB (4,831 insertions, 3,218 deletions), we find 13% of SV insertions and 15% of SV deletions have a greater offset.

By manually inspecting many SVs, MC appears to place SV breakpoints irrespective of small variants and does left-align against the reference. For example, a 2.1 kbp insertion was identified in all HGSVC haplotypes (AF = 100%) with breakpoint variation, but it is shifted by 43 bp in the MC graph. This variant inserted into a TE and had TE sequence at the breakpoint creating a tract of imperfect microhomology, and MC aligned 43 bp from the insertion sequence to the reference. As a result, two false SNPs are found in all haplotypes with the SV and may mislead downstream analyses. For example, SNPs linked to SVs do suggest mechanisms of SV formation (Beck et al., 2019; Carvalho and Lupski, 2016; Deem et al., 2011), although no point mutations were actually generated by this SV. (**Fig 4A**). This pattern was observed frequently in the MC callset.

**Figure 4:**
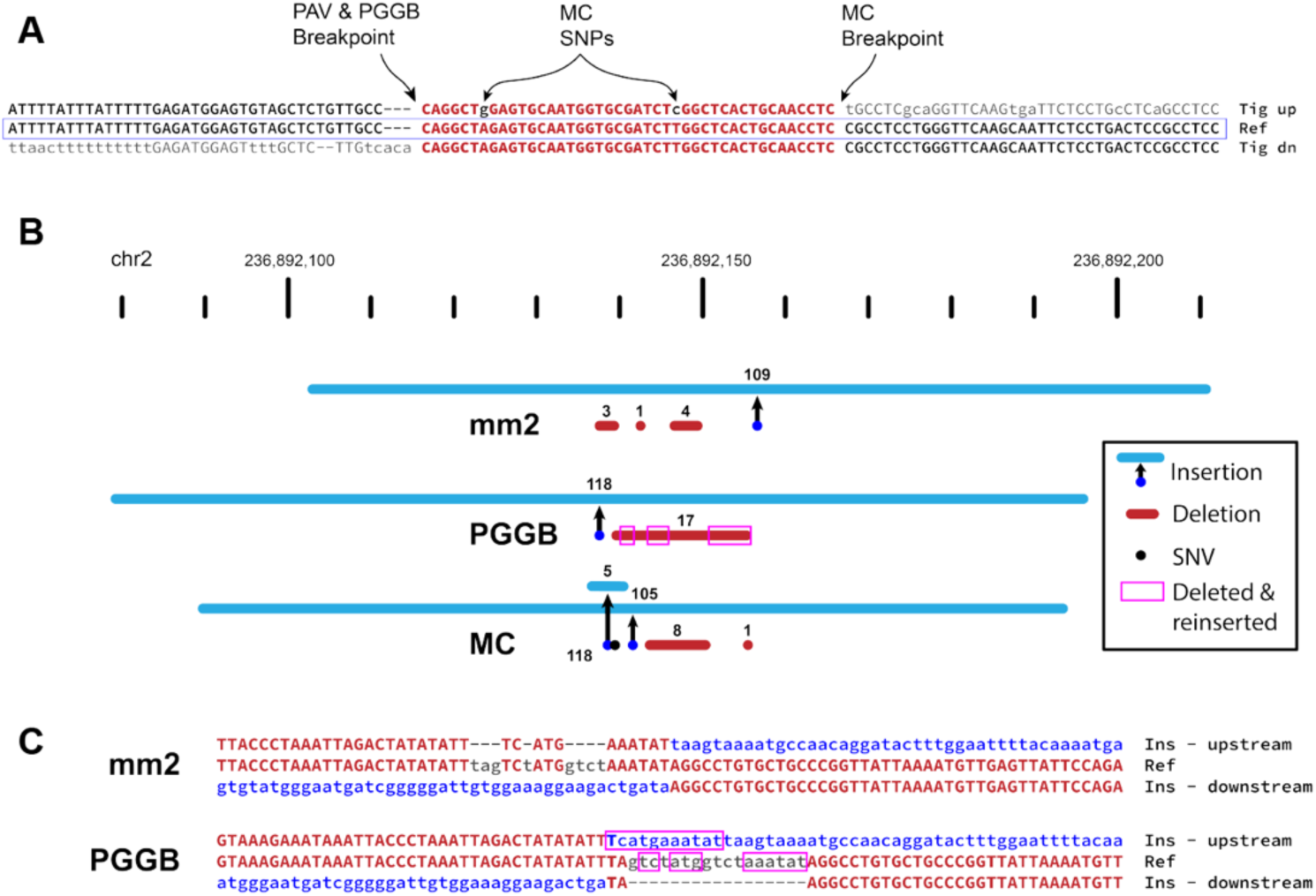
Pangenome graph breakpoints exhibit systematic bias. (A) A variant with a different breakpoint in the MC graph vs GRCh38 with the red portion showing the imperfect homology between breakpoint placements resulting in two SNPs called in the MC graph in all HPRC haplotypes (A>G and T>C). (**B**) An SV insertion (blue) paired with deleted bases (red) yields a 101 bp net gain by minimap2 (mm2, HGSVC callset), PGGB, and MC. minimap2 calls three small deletions near the insertion breakpoint. PGGB calls one larger deletion, but re-inserts deleted bases (magenta boxes) into the insertion call resulting in a larger SV insertion than minimap2. MC calls two insertions, two deletions, and a mismatch (black dot). (C) Alignments through breakpoints are shown for minimap2 and PGGB. Bases aligned to the reference are shown in red with matches in upper-case, inserted sequences are blue, and deleted bases are gray. The inserted sequence was not found in GRCh38, and likely represents the deletion of ancestral sequence, where the reference contains the derived (deleted) allele.

Many differences in the PGGB SVs are attributable to different breakpoint choices among largely equivalent representations. For example, a 162 bp VNTR expansion (27 bp motif) with one imperfect reference copy is inserted to the right of the reference copy rather than the left (**Fig S3**). More importantly, we find a distinct pattern of PGGB deleting and re-inserting the same bases when calling variants in loci without clean breakpoints. In one example, minimap2 represents a 101 bp net gain as a 109 bp insertion with three deletions totaling 8 bp, PGGB calls a 118 bp insertion with a single 17 bp deletion, and MC calls a 105 bp insertion, a 5 bp insertion, two deletions totaling 9 bp, and a SNP (**Fig 4B**). Further inspection of the breakpoints shows that 13 bases deleted by PGGB are re-inserted as part of the SV insertion (**Fig 4C**). This SV insertion sequence does not align to the human reference, but is present in the *Pan troglodytes* on Chromosome 2 and is also in other primate genomes. Therefore, the insertion and is likely ancestral and the deletion became the reference allele by chance. The minimap2 representation of this locus appears to be the most likely biological explanation for this event with small template switches within the replication fork, which is characteristic of some repair mechanisms (Carvalho et al., 2013), most notably MMBIR (Hastings et al., 2009). Given that the insertion is ancestral, the deletion and re-insertion of bases is less likely.

In addition to creating different representations of SVs, the area between breakpoints is often filled with small variants that are annotated differently across the haplotypes which may impact the interpretation of variants. These different breakpoints intersect coding sequences for 26 genes on average in MC and 5 genes on average in PGGB with additional discrepancies in UTRs and ncRNAs (**Table 2**). For example, we find a 180 bp insertion in *ESYT3* where minimap2 and PGGB place the breakpoint in an intron, but MC places it in an exon (**Fig 5**).

**Figure 5:**
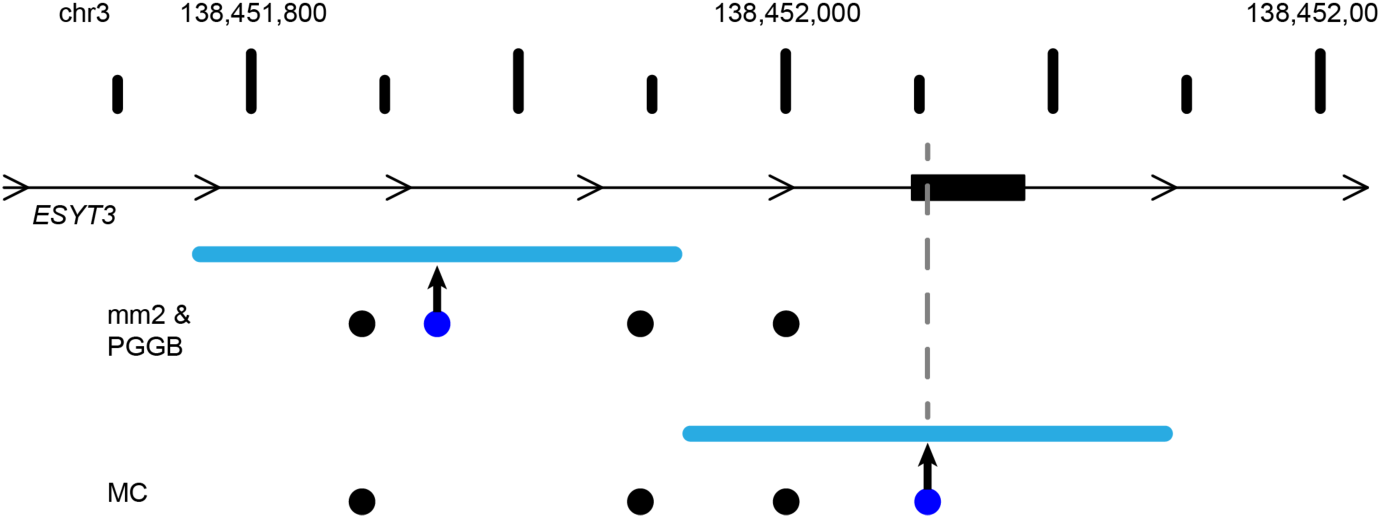
Breakpoint differences affect biological interpretation. A 180 bp insertion was called by minimap2 (mm2) and Pang at the same location within the intron of *ESYT3*, but Minigraph-Cactus (MC) placed the insertion 183 bp upstream inside an exon of *ESYT3*. SV insertions breakpoint locations are dark blue dots with the size of the SV shown as a light blue line. Point mutations are black dots. A gray line denotes the SV insertion location in the *ESYT3* exon.

**Table 2:** Consequences of differential breakpoints on RefSeq genes.

| Graph | SV Type | Sample | CDS | 5' UTR | 3' UTR | ncRNA |
| --- | --- | --- | --- | --- | --- | --- |
| mc | Insertions | HG03486 | 22 | 155 | 21 | 212 |
| mc | Insertions | HG02818 | 19 | 168 | 15 | 182 |
| mc | Deletions | HG03486 | 4 | 85 | 4 | 108 |
| mc | Deletions | HG02818 | 6 | 78 | 4 | 113 |
| pggb | Insertions | HG03486 | 12 | 53 | 8 | 74 |
| pggb | Insertions | HG02818 | 12 | 55 | 7 | 69 |
| pggb | Deletions | HG03486 | 3 | 41 | 2 | 47 |
| pggb | Deletions | HG02818 | 2 | 34 | 1 | 36 |

## Discussion

Advances in long-read sequencing coupled with new phased assembly and variant detection methods have increased the number of detectable SVs dramatically from less than 10,000 to more than 25,000 per diploid genome, and these advances continue to rival short-read technology by reducing costs, increasing availability, and improving read quality. In addition to detecting more SVs, long-reads also capture the full SV sequence, which is important for detailed analyses of non-reference sequences and has already proven to be transformative in mobile element characterization (Ebert *et al*., 2021; Ferraj *et al*., 2023).

Assemblies have improved breakpoint accuracy, however, systematic errors still exist where breakpoints span homologous regions. This effect is especially large for tandemly duplicated sequence and SV anchored in TEs. Since a majority of SVs have some form of homology around breakpoints (Ebert *et al*., 2021; Lam *et al*., 2010), the effect of differential breakpoints is potentially large even outside highly-repetitive sequences. However, modern aligners along with assembly-based SV detection consistently place SVs effectively reducing potential biases, but not eliminating them.

While errors in sequence and assembly do contribute to breakpoint differences, the most significant driver is the presence of small allelic changes near SV breakpoints, largely due to ancestral differences. This includes variation within insertions, which accumulate polymorphisms over generations. Short-reads are subject to known reference biases where distant haplotypes align less confidently to alternate reference alleles (Brandt *et al*., 2015; Degner *et al*., 2009). Although small polymorphisms are spanned by much longer flanking sequences with long reads and assemblies, this reference bias now manifests as differential breakpoints.

On a large scale, these small differences have little effect on callset quality because modern variant merging and comparison tools do allow for imprecise breakpoints, however, it does impact breakpoint annotations. This impedes precise mechanistic inferences since shifting a breakpoint changes microhomology annotations by an average of more than 20 bp and leads to a lack of polymorphisms flanking SVs. These polymorphisms can be signatures of the DNA repair causing the rearrangement (Beck *et al*., 2019; Carvalho and Lupski, 2016; Deem *et al*., 2011). As a result, callsets are still imprecise and incomplete, even within unique loci, despite being covered by long, contiguous, high-quality assemblies.

While pangenome graphs normalize SV loci across samples, additional developments are required to improve breakpoint precision. PGGB agrees with minimap2 for many SVs; some method tuning could potentially improve PGGB for SV breakpoints in unique loci, whereas more substantial improvements may be required for MC. As graph methods mature, they hold promise for calling variants at scale across divergent haplotypes. Importantly, rare and somatic variants will not be in graph references based on population samples, and calling variants against the closest reference path will face the many of the same challenges as methods based on linear references. Improving breakpoints in graph representations and linear references will ultimately increase the utility of pangenome references.

In this study, we investigate breakpoint disagreements in unique regions of the human genome where long reads and assemblies spanning rearrangements with megabases of flanking sequence and few assembly errors. Complex genomic loci are dense with repeats and breakpoint homology, and our results suggest that these loci present with larger breakpoint discrepancies. While these loci have added complexity from larger, more frequent, and more complex rearrangements as well as more collapsed reference loci (Sulovari et al., 2019; Vollger et al., 2022), making more rigorous methods for precise rearrangement breakpoints may help solve these regions more effectively.

Both simple and complex loci provide a rich opportunity for new methods to improve alignments, variant calling, and variant annotation. While current sequencing data captures these events with few errors, the limitations of current methods lead to systematic biases that affect the accuracy of variant calls and limit their utility for detailed downstream analyses. While long reads continue to gain in length and fidelity, the tools used to analyze them must keep pace.

## Methods

### Statistical analysis

Summary stats, such as mean and SD, were performed with Python numpy (v1.22.4) and statistical tests including Student’s t-test, Welch’s t-test, F-test, and Fisher’s exact test were carried out with scipy (1.9.3). All tests were two-tailed. F-tests were used to determine if a Student’s t-test was carried out (F-test *p*-value ≥ 0.01) or a Welch’s t-test (F-test *p*-value < 0.01).

*P*-values less than 1.0×10^-100^ are reported as p < 1.0×10^-100^. Extremely low *p*-values less than the smallest floating point value Python can represent (∼ 1×10^-308^ ± 1×10^-15^ on our system) are also reported as *p* < 1×10^-100^ in this manuscript.

#### Microhomology

The number of unique breakpoints was compared to the number of unique microhomology calls per merged variant. Neither the number of unique breakpoint locations and unique microhomology calls model a normal distribution (p < 1×10^-100^, scipy.stats.normaltest based on D’Agostino and Pearson’s test), so we computed correlation based on Spearman rank-order correlation coefficient.

### Genome reference

We use the hg38-NoALT reference published with the HGSVC callset (Ebert *et al*., 2021) (ftp://ftp.1000genomes.ebi.ac.uk/vol1/ftp/data_collections/HGSVC2/technical/reference/2020051 3_hg38_NoALT/). This reference is the full primary assembly of the human genome build 38 (GRCh38/hg38) including unplaced and unlocalized contigs, but it does not include patches, alternates, or decoys.

### Ebert callset

We acquired the version 2 (Freeze 4) merged callset from HGSVC (Ebert *et al*., 2021) (. We retained the same 32 population samples excluding the trio children used in the HGSVC publication. Frequencies and allele counts were adjusted to exclude child samples in the merged callset. We removed variants on unplaced and unlocalized contigs of the reference including only variants on primary chromosome scaffolds. A merging bug in SV-Pop allowed for some long-range intersects, and we removed these merged variants. To accomplish this filter this, we required either (a) the maximum offset is less than or equal to the merged SV length or (b) the maximum offset difference was less than 400 bp (200 bp in either direction) and the maximum SV length difference was not greater than 50% of the maximum SV length. These parameters mirror the expected results from the merging process without the long-range bug.

We obtained the Tandem Repeats Finder (TRF) (Benson, 1999) and RepeatMasker (Smit, 2013-2015) annotations from the UCSC Genome Browser (retrieved 2023-01-27, tracks "simpleRepeat" and "rmsk", respectively) (Kent et al., 2002). From TRF, we used all loci. From RMSK, we used all loci annotated as "Low_complexity" or "Simple_repeat". RMSK and TRF records within 200 bp were merged with BedTools merge (v2.30.0) (Quinlan and Hall, 2010) and a 200 bp flank was added to all merged regions to each and intersected variants with both TRF and RMSK. Insertions were marked as tandem repeats if their insertion point was within a padded repeat region. Deletions were marked as tandem repeats if either reference breakpoint was within a 200 bp padded repeat region. Intersects were done with BedTools intersect (v2.30.0) (Quinlan and Hall, 2010).

Segmental duplications were annotated using the same process as tandem repeats. The segmental duplication track was retrieved from the UCSC genome browser (2023-01-28, track " genomicSuperDups"). Regions were merged within 200 bp and a 200 bp flank was added, both operations with BedTools. Insertion and deletion breakpoints were intersected with the merged and padded SD regions in the same way as repeats.

Distance between variant breakpoints is defined as before in SV-Pop (https://github.com/EichlerLab/svpop) as used by HGSVC for merging: min([start offset, end offset]) where "start offset" is the distance between the variant start positions and "end offset" is the distance between the variant end positions, which may be different if the variant is a deletion (Ebert *et al*., 2021).

Pairwise comparisons were done by selecting all combinations of 64 haplotypes among the 32 samples (2,016 combinations of 2 haplotypes from a pool of 64). We obtained the original locations for each variant in all 64 haplotypes by tracing the merged call back through the sample to the original PAV call for each.

### TEMR annotations

We labeled SVs as TEMRs if TE annotations at SV breakpoints was consistent with a rearrangement mediated by TE homology. For reference repeats, we obtained the UCSC RepeatMasker (RMSK) track (database: rmsk.txt.gz) for hg38 (retrieved 2023-01-03). We retained only records with repeat class "LINE", "SINE", or "LTR" and with a minimum size of 100 bp. For deletions, we intersected the reference locations for each event independently (i.e. upstream breakpoint location and downstream breakpoint location) and annotated deletions as TEMRs if (a) both breakpoints intersected a TE annotation of the same type (e.g. Alu, ERV1, ERVK, L1, L2, etc), and (b) each side of the breakpoint intersected a different TE (i.e. distinct TE events). For SV insertions, we intersected the reference breakpoint with the RMSK track. We additionally obtained RepeatMasker annotations run on the merged callset by HGSVC (Ebert *et al*., 2021) and selected repeat annotations within 10 bp of each end of the insertion. We annotated insertions as TEs if (a) RMSK annotations at each end of the inserted sequence and at the reference breakpoint were all the same TE type, and (b) RMSK annotation at each end of the insertion sequence were not the same TE (i.e. distinct TE events). Breakpoint intersections with the RMSK track were performed with BedTools intersect (v2.30.0).

### Tandem duplication identification

Insertion sequences in unique loci (excluding annotated SDs and TRs) were re-mapped to the reference with BLAST (v2.13.0) with parameters "-word_size 16 -perc_identity 95" against a BLAST database constructed from hg38-NoALT. We compiled a list of filter regions by including all TRs and RepeatMasker (RMSK) annotations with a score of 50 or greater from the UCSC browser and merging records with BedTools merge (v2.30.0). BLAST alignments were discarded if 50% or more of the alignment record intersected the TR and RMSK filter. We further filtered BLAST hits to include only records that mapped within 10% of the SV length from the insertion site or deletion breakpoints (e.g. 1 kbp INS, 100 bp window around the insertion site) with a minimum of 100 bp for small SVs. For deletions, we removed the deletion sequence alignment (i.e. remapping produces an alignment over the deletion). Alignments less than 30 bp were also excluded. Some redundant overlapping alignments remained and appeared to be driven by small TRs that were not in the reference, which were removed by keeping only the longest record if records overlapped by 80% or more. The same 80% overlap was performed in both reference space using aligned reference coordinates and in SV sequence space using coordinates from the SV sequence (i.e. the first base of the SV sequence is position 0). We selected SVs where the total number of aligned bases on each side of the breakpoint was within 90% of the total SV size and ensuring records with large gaps spanning more bases than were aligned did not contribute to the SV size calculation. We did not select records that had the expected alignment pattern (i.e. upstream SV sequence mapping downstream of the SV breakpoint and downstream SV sequence mapping upstream of the SV), although all the records left after the filtering process did exhibit this pattern.

### Small variants around breakpoints

Our goal was to identify small differences between haplotypes that causes variant breakpoints to be placed differently. For each haplotype pair, we selected SV insertions and deletions with breakpoints placed at different sites and with breakpoints in unique loci (not TR or SD). We extracted the haplotype sequence from around the assembly including a 50 bp flank on each side and we extended one end appropriately to add flank so that in the absence of other small variants, both sequences should start on the same base.

The sequences were aligned so that the right-most variant was the reference and the left-most variant was the query, although either order should produce similar results. Sequences were aligned with the "swalign" Python package (v0.3.7) using a global alignment and with match,

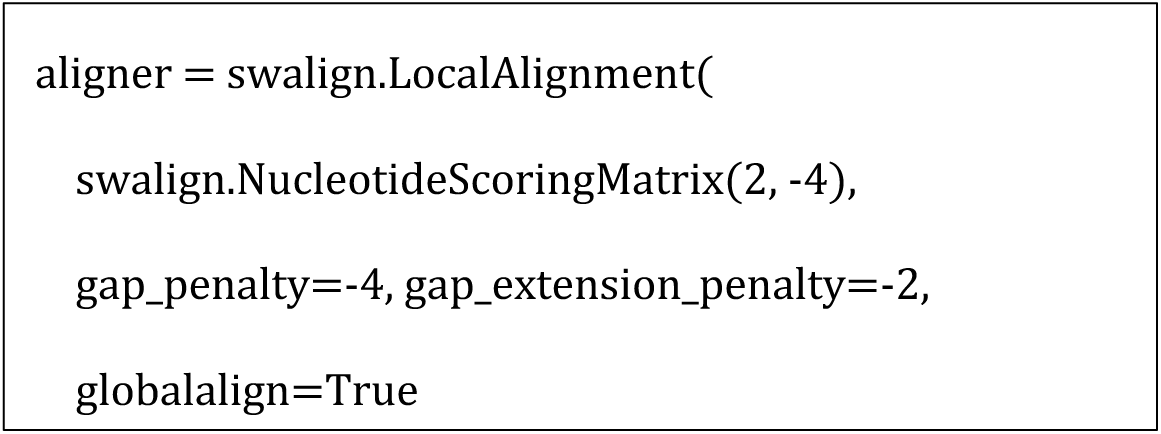

mismatch, and gap scores from the minimap2 (short-gap scores from the default double-affine parameters):

Alignment align ("M" CIGAR operations) records were transformed to match/mismatch ("=" and "X" CIGAR operations), and using the known flanks added to each, we assigned variants to left flank, left breakpoint (intersecting the breakpoint), differential region, right breakpoint (intersecting the breakpoint), and right flank along with their relative position in each category.

### Microhomology

Microhomology is the span of perfectly matching bases at each end of a breakpoint, for example, the perfect homology at sites of ectopic recombination (i.e., NAHR), homology directed repair, replication-based repair, or alt-EJ. We measured homology at breakpoints using an algorithm in PAV and previously validated as part of a TEMR project (Balachandran *et al*., 2022) where the region upstream of an SV sequence is matched with the downstream reference or contig, and the region downstream of the SV sequence is matched with the upstream reference or contig. To compare haplotypes more consistently, we computed SV homology for insertions against the upstream and downstream contig where the SV was called, and against the reference for deletions. We excluded all TD variants from homology because estimating breakpoint homology using this method counts whole TD copies as homologous.

### Graph genome comparisons

Variants against GRCh38 for PGGB and MC graphs were obtained from the decomposed VCFs published by the HPRC (Liao *et al*., 2023) (see "Resource table" in Methods for URLs). Variants were extracted for each sample using SV-Pop (Ebert *et al*., 2021). Differences were manually investigated using the UCSC browser and custom browser tracks for HGSVC and HPRC variants.

### Resource table

| Resource | Source | Identifier / version |
| --- | --- | --- |
| Merged Ebert callset | Ebert *et al.* (2021)<br>https://ftp.1000genomes.ebi.ac.uk/vol1/ftp/data_collections/HGSVC2/release/v2.0/integrated_callset/ | v2.0 (Freeze4) |
| Python |  | 3.9.15 |
| Numpy |  | 1.22.4 |
| Scipy |  | 1.9.3 |
| BedTools |  | 2.30.0 |
| BLAST |  | 2.13.0 |
| swalign | Python package | 0.3.7 |
| HPRC PGGB decomposed VCF | https://s3-us-west-2.amazonaws.com/human-pangenomics/pangenomes/freeze/freeze1/pggb/vcfs/hprc-v1.0-pggb.grch38.vcfbub.a100k.wave.vcf.gz |  |
| HPRC MC decomposed VCF | https://s3-us-west-2.amazonaws.com/human-pangenomics/pangenomes/freeze/freeze1/minigraph-cactus/hprc-v1.0-mc.grch38.vcfbub.a100k.wave.vcf.gz |  |

## Acknowledgments

P.A.A and C.R.B were supported by NIH NIGMS R35GM133600 and NIH NCI P30CA034196. The Human Genome Structural Variation Consortium (HGSVC) provided published data, support, and feedback, and the HGSVC was supported by NIH NHGRI U24HG007497. Thanks to Parithi Balachandran for helping to check duplication detection tools and SV breakpoints. Thanks to Evan E. Eichler for providing feedback on graph genome comparisons.

## Supplemental Material

**Figure S1:**
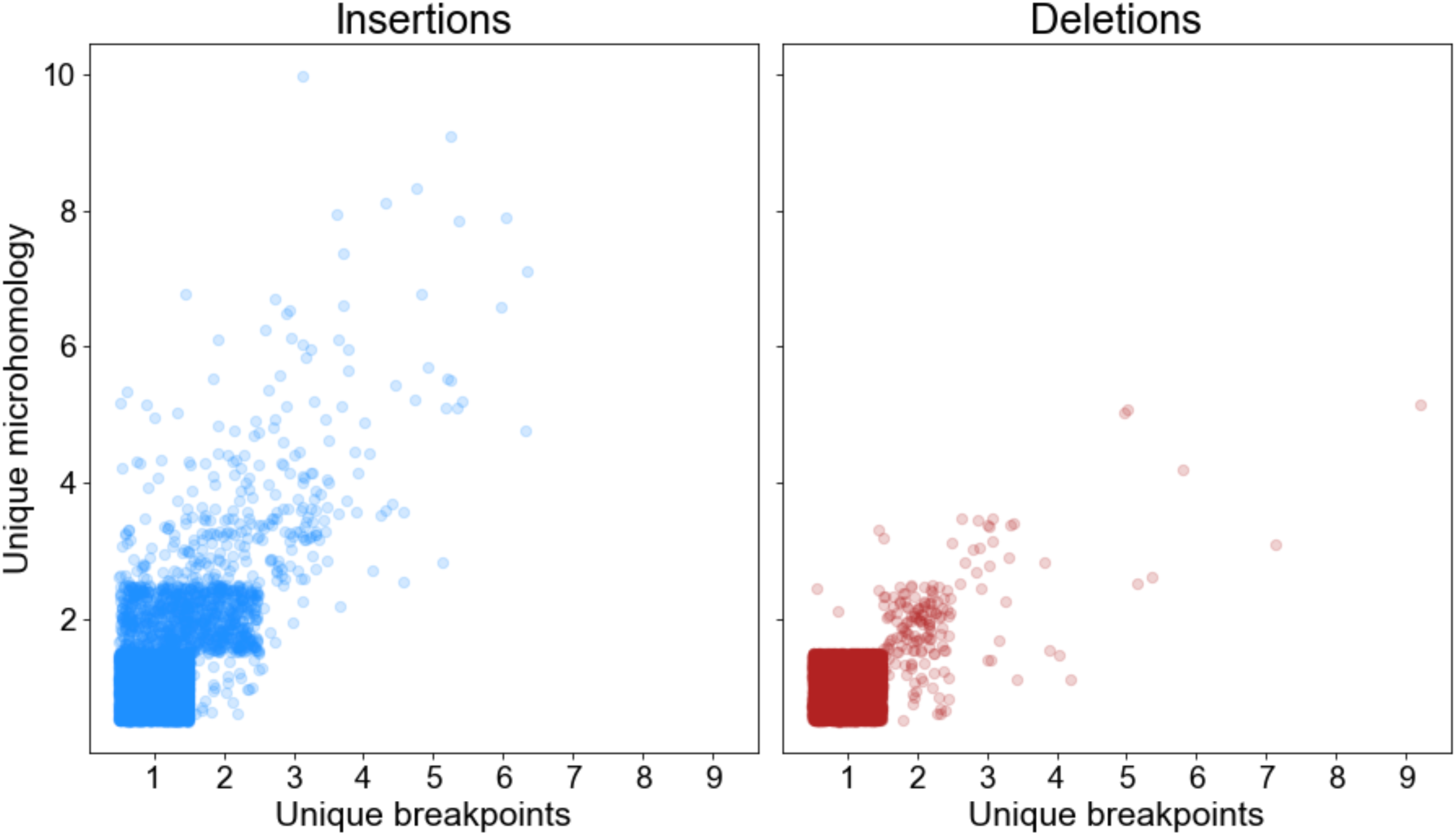
Breakpoint changes alter microhomology. The number of unique breakpoints (horizontal axis) a variant has across haplotypes has a dramatic impact on the number of unique microhomology annotations (violet line: best-fit least-squares regression line). Transparency and jittering (± 0.5) separates points falling on integers.

**Figure S2:**
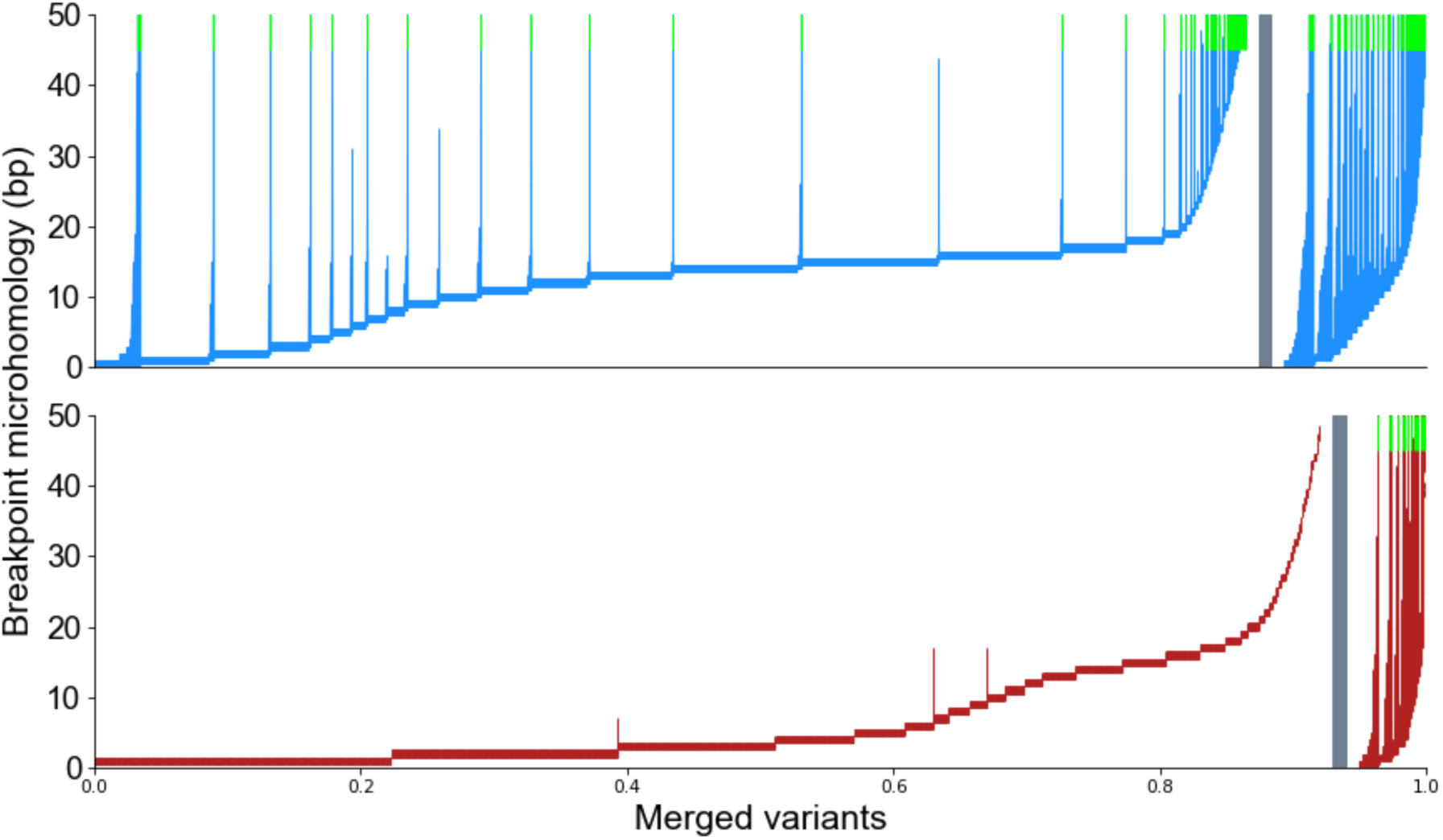
Breakpoint placement leads to large microhomology differences. For each merged SVs (insertions top/blue, deletions bottom/red), vertical lines extend from the minimum microhomology to the maximum microhomology across haplotypes. A gray bar separates SVs with consistent breakpoints (left) from SVs called at different breakpoints across haplotypes (right). Green tips denote lines that extend past the top of the figure.

**Figure S3:**
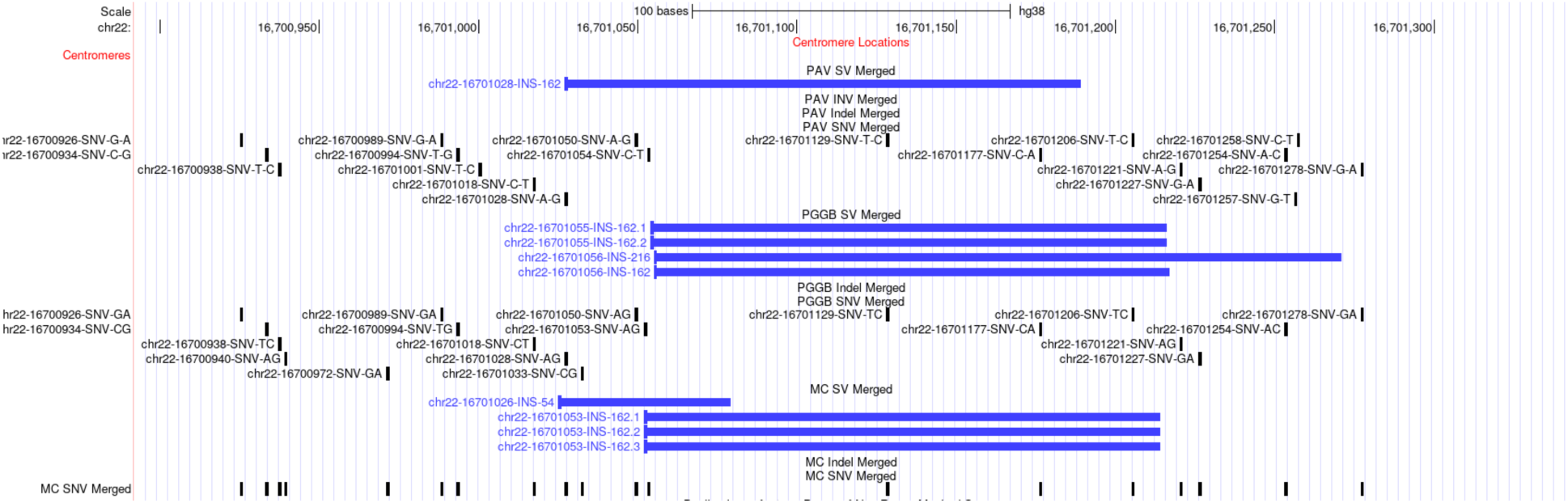
Ambiguous breakpoints for SVs in degenerate tandem repeats. The true breakpoint for this 162 bp expansion is difficult to identify even though tandem repeats in this locus were too diverged or too small to yield a tandem annotation. Despite this divergence, breakpoints were still not consistently placed, and the optimal location is difficult to identify and all three methods chose different breakpoints.

**Table S1.**
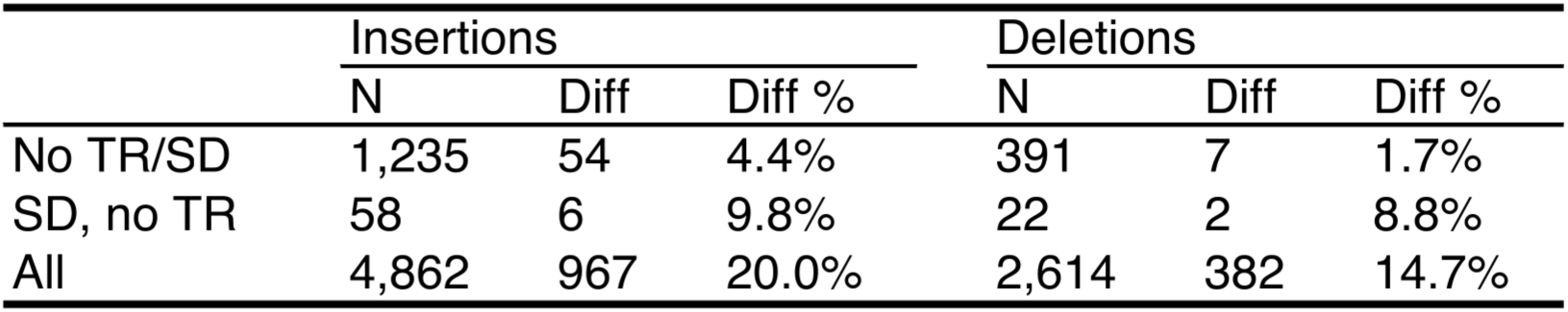
Offsets per haplotype pair. Average effect of shifted breakpoints on each pair of haplotypes (n = 2,016 combinations of 64 haplotypes). TR: Tandem Repeat, SD: Segmental Duplication, N: Number of variants, Diff: Variants with different offsets in the pair of haplotypes.

|  | Insertions |  |  | Deletions |  |  |
| --- | --- | --- | --- | --- | --- | --- |
|  | N | Diff | Diff % | N | Diff | Diff % |
| No TR/SD | 1,235 | 54 | 4.4% | 391 | 7 | 1.7% |
| SD, no TR | 58 | 6 | 9.8% | 22 | 2 | 8.8% |
| All | 4,862 | 967 | 20.0% | 2,614 | 382 | 14.7% |

